# Simultaneous single-cell profiling of the transcriptome and proteome

**DOI:** 10.64898/2026.05.14.724921

**Authors:** Xuling Xu, Monica Pia Caggiano, Malcolm L. Wells, Guizhi Sun, Sue Mei Lim, Dylan H. Multari, Scott A. Blundell, Nicolas Hartel, Rosa Viner, Jose M. Polo, Ralf Schittenhelm, Alex de Marco

## Abstract

Transcriptomic and proteomic measurements from the same single cell provide complementary information that cannot be inferred from either modality alone, yet methods for the parallel recovery of both analyte classes from a single-cell lysate remain limited. Here, we describe a workflow in which individual cells are isolated by automated dispensing into a minimal, MS-compatible lysis volume, followed by sequential mRNA capture and protein supernatant recovery, prior to independent downstream processing. The method is compatible with standard library preparation and data-independent acquisition proteomics pipelines and requires no dedicated instrumentation beyond a single-cell dispensing platform.

We evaluated workflow performance on 67 single cells across 3 iBlastoids. Transcriptomic sequencing detected a median of 5375 genes per cell, and proteomic analysis identified a median of 2123 protein groups per cell across two mass spectrometry platforms. Compared with a standalone single-cell proteomics protocol, incorporating the mRNA extraction step reduced median proteomic depth by approximately 11% (median 1,965 vs. 2,204 protein groups per cell), while mean per-cell identification remained comparable across workflows (1,790 vs. 1,775 protein groups per cell). Direct comparison of paired transcript and protein abundance yielded a median Spearman correlation of ρ ≈ 0.38; after correction for detection depth, the partial correlation was 0.067.

## Introduction

Cellular phenotypes are determined by the interplay among multiple molecular layers, with the proteome serving as the primary functional output. While transcript abundance provides a broad view of gene regulatory activity, protein levels are subject to extensive post-transcriptional control, including translational regulation, differential protein stability, and post-translational modification, such that mRNA abundance is an imperfect proxy for protein abundance ^1-4^. Direct measurement of both modalities from the same cell is therefore required to examine the mRNA-protein relationships without confounding contributions from cell-to-cell variability inherent to unpaired experimental designs ^5^.

Single-cell RNA sequencing has achieved broad adoption due to its scalability and sensitivity, enabling transcriptome-wide measurements across thousands of cells per experiment ^6-8^. Advances in mass spectrometry-based single-cell proteomics (scProteomics) have recently enabled the detection of a few thousand proteins from individual cells, facilitated by miniaturized sample-preparation platforms, improved instrument sensitivity, and data-independent acquisition strategies^9-12^.

Early approaches to single-cell multi-omics combined transcriptomics with targeted protein quantification using antibody-derived tags. CITE-seq ^13^ and related strategies enable simultaneous measurement of the transcriptome and a predefined set of surface proteins within the same cell and have been widely applied for cell-type characterization ^13, 14^. However, these methods are constrained by antibody availability and specificity, typically cover only tens to a few hundred protein targets, and are limited to epitopes accessible for antibody labeling, thereby excluding intracellular proteins, post-translational modifications, and protein isoforms. Mass spectrometry offers an unbiased alternative that can detect proteins without prior selection and captures a substantially broader fraction of the proteome, including nuclear and cytoplasmic proteins, modified peptides, and protein isoforms. Integrating MS-based proteomics with transcriptomics at the single-cell level requires that both analyte classes be recovered from the same cell. This constraint places significant demands on sample handling, reagent compatibility, and minimization of material loss during preparation.

Two broad strategies can be explored to address this challenge. Physical splitting of single-cell lysates into separate fractions for independent omics analyses has been demonstrated as a strategy for paired measurements ^5^. Implementation using nanoliter-scale droplet platforms has enabled global proteomic and transcriptomic profiling from the same cell ^15^. Alternatively, sequential biochemical fractionation, in which one analyte class is selectively captured prior to recovery of the other from the same volume, offers a route to parallel measurement without requiring physical division of the sample. Oligo(dT) magnetic bead capture of polyadenylated RNA is a well-established biochemical separation step compatible with downstream transcriptomic library preparation and has been incorporated into bulk multi-omics workflows ^16^. Whether this strategy can be applied at the single-cell scale while preserving sufficient protein material for MS-based proteomics has not been systematically evaluated.

Here, we describe an integrated workflow that combines automated single-cell dispensing into an MS-compatible lysis buffer, followed by sequential poly(A) mRNA capture and protein supernatant recovery, enabling independent downstream processing. We benchmark workflow performance across 67 single cells on two mass spectrometry platforms and characterize the effect of the mRNA extraction step on proteomic depth. As a proof-of-concept, we apply the workflow to human iBlastoids, which are synthetic three-dimensional cellular structures obtained by cellular reprogramming of fibroblasts that recapitulate the major cell lineages of the early human blastocyst, including epiblast, trophectoderm, and primitive endoderm ^17^. We show that paired transcriptomic and proteomic measurements from the same cells reveal coordinated and divergent molecular signatures across lineages that cannot be fully resolved from either modality alone.

## Results and Discussion

### An integrated workflow for parallel single-cell transcriptomics and proteomics

Single cells were isolated by automated dispensing using a cellenONE platform, selecting cells based on diameter (14-32 µm) and elongation (≤2) to exclude debris, doublets, and apoptotic cells (Supplementary Fig. 1). Each cell was dispensed into a pre-filled well of a 384-well low protein-binding plate containing 2 µL of MS-compatible lysis buffer comprising 0.4% DDM, 200 mM TEAB (pH 8.0), 3 ng trypsin, and 3 ng LysC. Following lysis, oligo(dT) magnetic beads were added to capture polyadenylated RNA, and the protein-containing supernatant was recovered by magnetic separation for downstream proteomic processing. Captured mRNA was independently eluted and processed for library preparation. The two downstream steps, transcriptomic library preparation and proteomic sample digestion, were performed on the same cellular lysate without physically splitting the sample (Fig. 1A).

**Figure 1.**
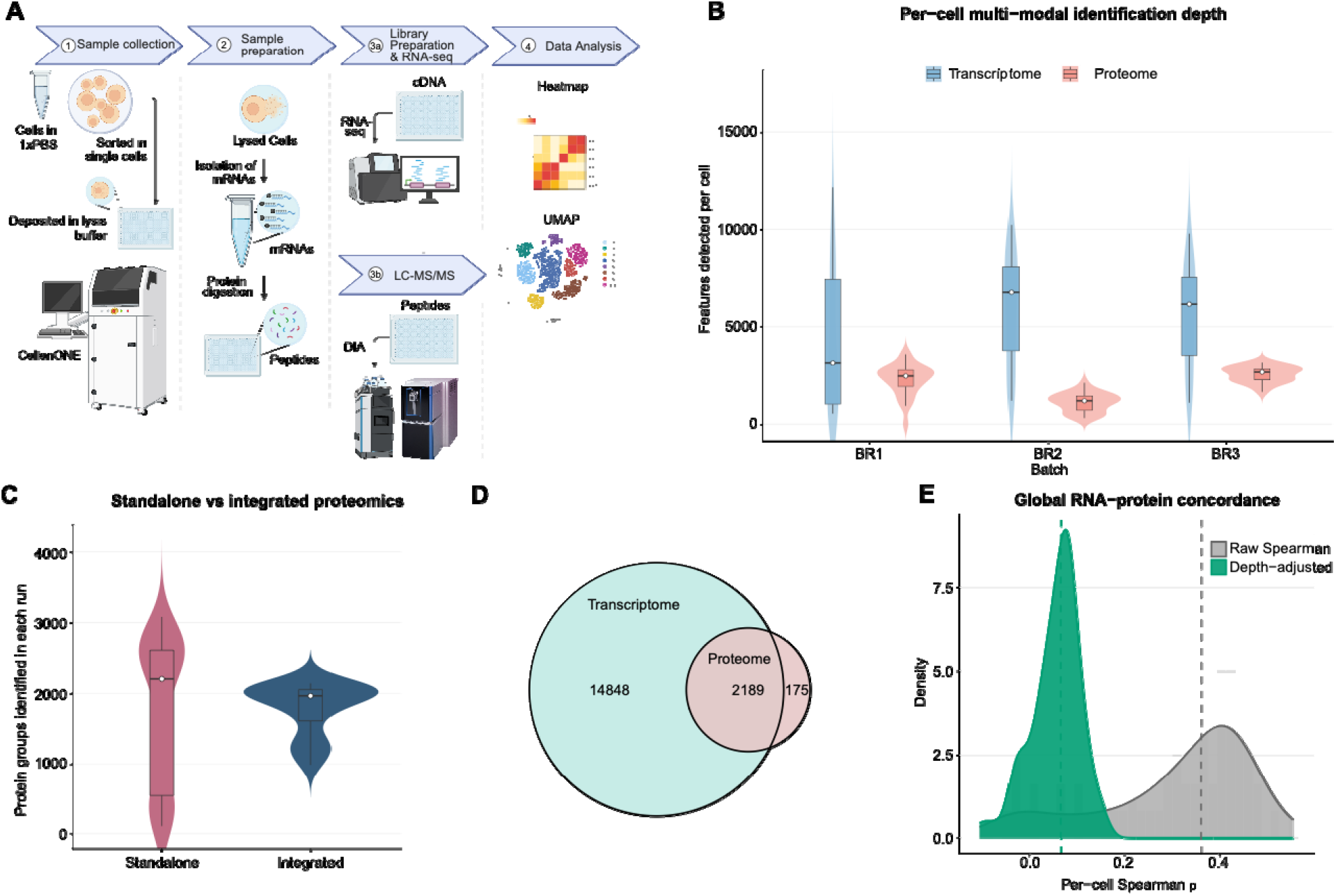
Workflow benchmarking and multi-modal feature detection. Panel A: Schematic of the workflow; Panel B: Per-cell multi-modal identification depth across all cells, grouped by batch (BR1, BR2, BR3); Panel C: Standalone single cell proteomics vs. integrated single cell transcriptomics and proteomics comparison, ranked by protein ID count; Panel D: Venn diagram of transcriptome/proteome feature overlap; Panel E: Global distribution of per-cell Spearman ρ with depth-corrected overlay.

### Benchmarking transcriptomic and proteomic depth

Across 67 single cells processed through the integrated workflow in three independent sample batches (BR1, BR2, BR3), transcriptomic sequencing detected more than 17,000 genes at the dataset level, with a median of 5375 genes detected per cell (Fig. 1B). Proteomic analysis identified more than 4,000 protein groups at the dataset level, with per-cell yields ranging from approximately 1,000 to 3,000 protein groups depending on the mass spectrometry platform used. Samples analyzed on the Orbitrap Astral platform (BR1 and BR3) yielded a median of 2,534 protein groups per cell, while samples analyzed on the Orbitrap Exploris 480 (BR2) yielded a median of approximately 1196 protein groups per cell. The higher identification depth on the Astral platform is consistent with its higher scanning speed and sensitivity relative to the Exploris 480 and reflects instrument-dependent rather than protocol-dependent variation.

At the mean level, per-cell protein group identification was comparable between workflows (integrated: ∼1,790; standalone: ∼1,775). At the median level, the integrated workflow identified 1,965 protein groups per cell compared with 2,204 for the standalone protocol, representing an 11% reduction. Per-cell protein-group counts clustered primarily by sample batch rather than by workflow, indicating that the dominant source of variance is biological/sample input rather than the integrated versus standalone protocol (Fig. 1C, Supplementary Fig. 2). The broader distribution observed in the integrated workflow is consistent with this batch effect and may additionally reflect variable mRNA capture efficiency or material loss during magnetic separation. Of the protein groups identified across the two protocols, approximately 54% were detected by both, while 37% were unique to the standalone condition and 9% unique to the integrated workflow (Supplementary Fig. 2). Despite this reduction, per-cell identification counts in the integrated workflow remain within the range reported in current single-cell proteomics benchmarks ^9, 10, 12^, indicating that the sequential extraction design preserves sufficient proteomic depth for downstream analysis.

### Non-redundant feature detection across modalities

Across the full dataset, 2,189 molecular features were detected at both the mRNA and protein level, while 14,848 features were detected exclusively in the transcriptomic data, and 175 protein groups had no corresponding mRNA detection (Fig. 1D). The larger RNA-exclusive set reflects the broader dynamic range and sensitivity of sequencing relative to mass spectrometry. The protein-exclusive detections likely arise from post-transcriptional buffering, transcript degradation prior to library preparation, or stochastic dropout in the RNA modality. These distributions confirm that the two modalities provide substantively non-redundant molecular information from the same cell.

### RNA-protein correlation and the contribution of the technical depth

A direct comparison of paired transcript and protein abundance across single cells yielded a global median Spearman correlation of ρ ≈ 0.38 (Fig. 1E). Per-cell correlation values increased with both RNA detection depth and protein identification count; cells with greater coverage in both modalities showed higher apparent RNA-protein concordance. This pattern is expected from a cell-level sampling effect: cells that are more deeply sampled in both modalities have a broader dynamic range and more co-detected features, producing more stable and inflated correlation estimates, independent of the underlying biology. To isolate the cell-intrinsic biological signal from this sampling-depth covariation, expression values were residualized against per-cell detection counts for each modality prior to computing correlations. After this correction, the depth-adjusted median correlation shifted to 0.067 (Fig. 1E), indicating that a substantial fraction of the raw RNA–protein concordance reflects shared sampling efficiency across modalities rather than coordinated gene regulation. A one-sample Wilcoxon test confirmed that depth-corrected per-cell ρ values were significantly greater than zero across the full dataset (p < 0.001) and within each batch independently (B1: p = 1.6e^−^□; B2: p = 2.0e^−^□; B3: p = 5.2e^−^□), with no significant difference in residual ρ across batches (Kruskal-Wallis, p = 0.14), indicating that the result is reproducible and not attributable to instrument-specific technical variation. The persistence of a small but significant residual correlation (per-batch medians: 0.073, 0.041, 0.076) suggests that a detectable component of RNA–protein concordance reflects genuine co-regulation, consistent with the subset of genes subject to tight translational coupling ^1, 15, 18, 19^. Nevertheless, the magnitude of the corrected correlation remains substantially lower than bulk-level estimates, consistent with prior reports documenting modest transcript– protein concordance at the single-cell level, and underscores the extent to which post-transcriptional regulation and differential protein stability decouple the two molecular layers in individual cells.

### Cell population structure resolved by paired multi-modal profiling

As a proof of concept, the workflow was applied to 67 single cells derived from human iBlastoid models, which comprise multiple distinct cell populations. Unsupervised graph-based clustering of the transcriptomic data identified two primary cell clusters, which were visualized by UMAP (Fig. 2A, left). Further projection onto a published single-cell reference atlas ^17^ using anchor-based label transfer, assigning cells to trophectoderm-like (TE), epiblast-like (EPI), primitive endoderm-like (PE), and intermediate states (Fig. 2A, right), consistent with the known lineage composition of the iBlastoid model ^17^.

iBlastoids contain epiblast (EPI)- and trophectoderm (TE)-like populations that can be distinguished by their canonical lineage markers. We therefore compared cells with high SOX2 expression (a canonical EPI marker) against cells with high GATA2 expression (a canonical TE marker), and identified distinct transcriptomic and proteomic signatures in each population (Fig. 2B, Supplementary Fig. 3). Additional canonical lineage markers (OCT4, NANOG, CDX2) showed concordant expression patterns and are presented in the Supplementary Data; SOX2 and GATA2 are highlighted here as representative EPI and TE markers. GATA2-high cells were uniformly classified as TE by reference projection, whereas SOX2-high cells were not exclusively annotated as EPI (likely because global transcriptomic profiles weight TE identity in this dataset, even in cells with elevated SOX2). At the RNA level, SOX2-high cells showed enrichment for gene ontology terms related to ciliogenesis, while GATA2-high cells were enriched for Prostaglandin secretion. At the protein level, SOX2-high cells showed enrichment for response to transforming growth factor beta and centrosome localization, and GATA2-high cells for establishment of organelle localization (Fig. 2C-D). The partial overlap between RNA and protein GO signatures within each population illustrates the complementary resolution afforded by paired measurement: pathway-level concordance is detectable while modality-specific features remain distinct.

**Figure 2.**
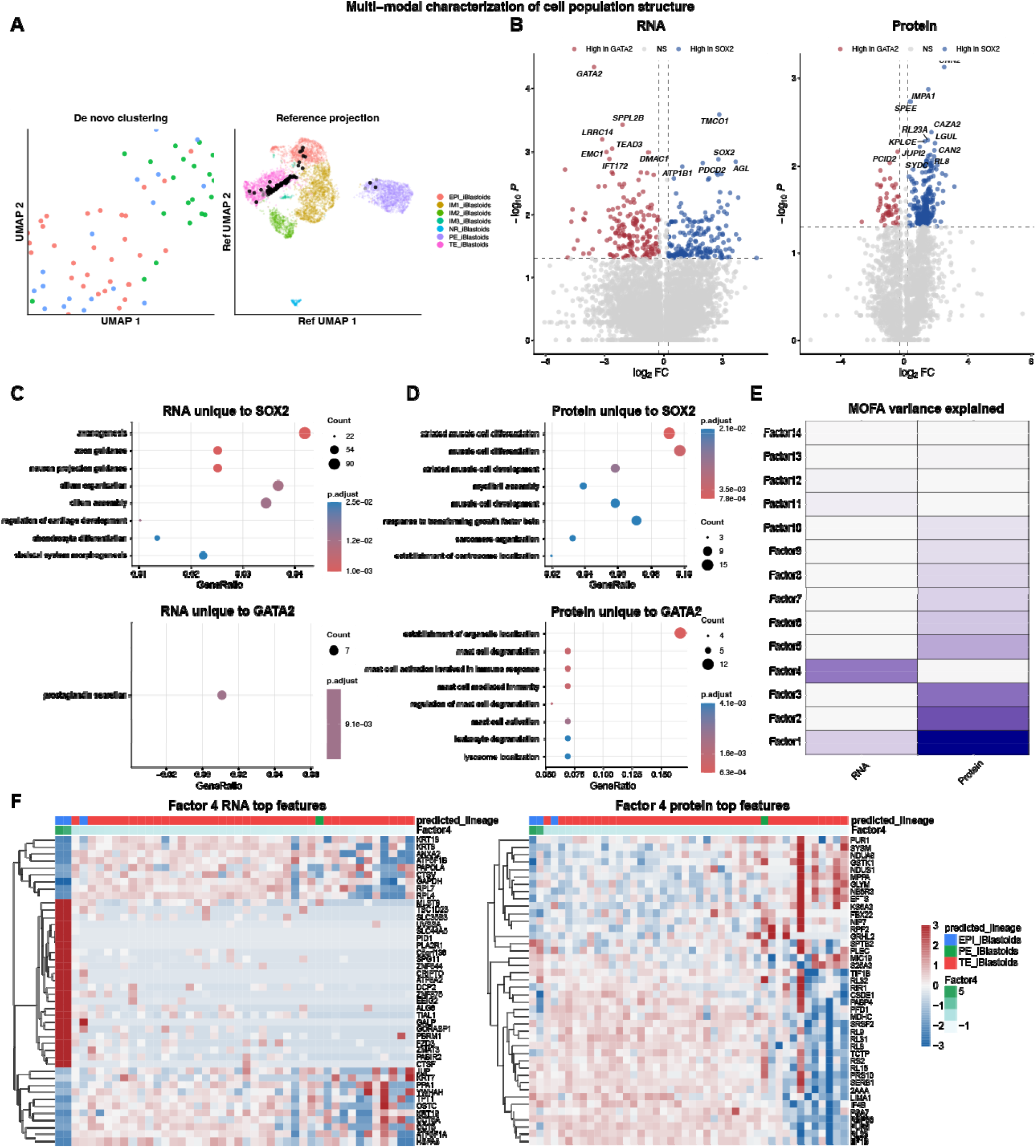
Multi-modal characterization of cell population structure. Panel A: UMAP de novo clustering and reference projection; Panel B: Volcano plots RNA and protein, SOX2 vs GATA2; Panel C: GO enrichment dot plots for RNA unique features; Panel D: GO enrichment dot plots for protein unique features; Panel E: MOFA variance explained; Panel F: MOFA feature heatmaps for Factor 4 (lineage factor).

### Multi-modal factor analysis identifies biological and technical axes of variation

To systematically decompose sources of variation across both modalities, we applied Multi-Omics Factor Analysis (MOFA2) to the integrated dataset, training a model across 10 factors (Fig. 2E). Factors were separated into three interpretable categories: those capturing technical depth variation, those reflecting coordinated biological programs across both modalities, and those dominated by proteome-specific signals (Supplementary Table 1).

Factors 1 and 3 showed strong correlations with RNA and protein detection rates, respectively, consistent with modality-specific technical variation in sampling depth rather than biological signal (Supplementary Fig. 4, Supplementary Fig. 5). Factor 2 was dominated by the proteomic modality and captured variation in ribosomal and ribonucleoprotein complex components, with GO enrichment for membraneless organelle assembly and stress granule formation, suggesting heterogeneity in biosynthetic and post-transcriptional regulatory capacity (Supplementary Fig. 6, Supplementary Fig. 7). Factor 4 showed the strongest association with cell lineage identity: a Kruskal-Wallis test indicated significant differences in factor scores across lineages (p = 0.0147), with pairwise testing confirming significant separation between TE-like and EPI-like populations (BH-adjusted p = 0.00097). Lineage identity explained approximately 63% of variance in Factor 4, while correlations with RNA and protein detection rates were weak (ρ ≈ 0.11 and 0.07, respectively), confirming a biological rather than technical origin (Fig. 2E) (Supplementary Fig. 8, Supplementary Fig. 9). The top RNA and protein features loading on Factor 4 showed coherent expression gradients across cells ordered by factor score, with distinct patterns in TE-like versus EPI-like populations (Fig. 2F, Supplementary Fig. 10).

The ability to identify a lineage-associated factor with significant cross-modal coupling demonstrates that it is possible to resolve both the transcriptome and the proteome from the same single cell, with sufficient depth and without reliance on computational integration of unpaired datasets. Factor 4 captures a biological axis detectable in both molecular layers simultaneously, suggesting that the transcriptional and proteomic states that define lineage identity are sufficiently coupled to be recovered from the same cell. Such paired analyses make it possible to identify regulatory transitions in which RNA and protein signatures diverge. For example, cases where lineage identity is established or maintained at the protein level independently of transcript abundance ^5, 20^, or where post-transcriptional buffering decouples the two layers during cell-state transitions ^18^. These are questions that cannot be addressed by individual measurements or by cross-cell computational integration, in which cell-to-cell variability confounds the interpretation of modality-specific differences, and the time component cannot be unequivocally deconvolved. Direct paired measurement, as demonstrated here, provides the molecular resolution required to distinguish genuine post-transcriptional regulation from technical noise in single-cell multi-omics data.

## Methods

### iBlastoid generation

iBlastoids were generated using a modified version of the protocol described by Liu et al. (2021) in Nature ^17^. Briefly, primary human adult dermal fibroblasts (HDFs) derived from female donors were reprogrammed using the CytoTune-iPS 2.0 Sendai Reprogramming Kit (ThermoFisher Scientific). Sendai viral vectors encoding KOS, c-MYC, and KLF4 were applied at multiplicities of infection (MOI) of 5:5:6 KOS (KLF4, OCT4, SOX2), hc-MYC, and hKLF4, respectively. Reprogramming was carried out in fibroblast medium for 21 days. Day 21 intermediate cells were dissociated and seeded into Corning^®^ Elplasia^®^ 6-well plates (Corning^®^) at a density of 100 cells per microwell in modified human iBlastoid medium.

### iBlastoid collection, dissociation, and flow cytometry

Day 7 MD2 and MD4 iBlastoids were collected and processed under the following conditions.

For lysis-based downstream applications, Day 7 iBlastoids were pooled, washed once in phosphate-buffered saline (PBS), and resuspended in 100 mL lysis buffer. Samples were then transferred for downstream processing.

For additional processing, Day 7 iBlastoids were collected, washed once in PBS, and dissociated using TrypLE^™^ Express Enzyme (1X) (Thermo Fisher Scientific) at 37 °C for 3 mins with mechanical trituration to obtain single-cell suspensions. Cells were diluted in PBS, centrifuged at 300 × g for 5 min, washed once more in PBS, and resuspended in 200 mL PBS prior to downstream applications.

Unless otherwise stated, all samples were maintained on ice throughout processing, and all buffers were pre-chilled.

### Single-cell isolation

Harvested iBlastoid cells were washed twice in ice-cold 1xPBS and resuspended to 200–400 cells/µL. Single cells were isolated and dispensed into 384-well low-protein-binding Armadillo PCR plates (ThermoScientific AB2384) using a cellenONE instrument equipped with a medium-sized PDC, selecting cells with a diameter of 14-32 µm and an elongation ≤2 to exclude debris, doublets, and apoptotic cells. (24.2% isolation rate, 397 single particles detected, of which 96 passed the isolation criteria). Each well was pre-filled on ice with 2 µL of 2x MS-compatible lysis master mix containing 0.4% dodecylmaltoside (DDM) (Thermo Fisher Scientific - 89902),200 mM TEAB (pH 8.0) (Sigma - T7408-100ML), 0.3 ng/µL trypsin gold (Promega - V5280), and 0.3 ng/µL LysC (Promega-VA1170). The plate was placed into the pre-cooled (4°C) cellenONE prior to cell sorting. After sorting, the plate was sealed with a silicone lid (Axygen, cat. no. AX-AM-384-PCR-RD) and either processed immediately or stored at −80°C.

### Integrated mRNA and protein extraction

Cell lysates were transferred from the 384-well plate to 8-strip PCR tubes. 0.5 µL of Invitrogen Dynabeads Oligo(dT)25 magnetic beads (Thermo Fisher Scientific - 61002) per single cell were washed in 10 µL of binding buffer (20mM Tris-HCl pH7.5, 1.0M LiCl, 2mM EDTA) and resuspended in 2 µL of 2× lysis/binding buffer (200mM Tris-HCl pH7.5, 1.0M LiCl, 20mM EDTA, 10mM DTT), and added to the lysate. mRNA was allowed to anneal to the oligo(dT) sequences for 3-5 minutes at room temperature. The protein-containing supernatant was then recovered by magnetic separation and transferred back to the original wells of the 384-well plate for proteomic sample preparation. Beads were washed twice with 100µL of Washing Buffer A (10mM Tris-HCl pH7.5, 0.15M LiCl, 1mM EDTA and 0.1% LiDS) and twice with 100µL of Washing Buffer B (10mM Tris-HCl pH7.5, 0.15M LiCl, 1mM EDTA). mRNA was eluted by removing the beads from the magnetic separator and incubating in 10.5 µL of 10 mM Tris-HCl at 75-80°C for 2 minutes. Eluted mRNA was processed immediately for library preparation or stored at −80°C.

### Proteomic sample preparation and LC-MS analysis

The recovered protein supernatant was subjected to on-plate digestion at 50°C with 85% humidity for 2 hours inside the cellenONE instrument. Digestion was quenched with 1 µL of 1% acetic acid and supplemented with 2 µL of loading buffer (2% Acetonitrile, 0.1% TFA). Peptides were analyzed using a Vanquish Neo^™^ UHPLC system coupled to either an Orbitrap Exploris 480^™^ or an Orbitrap Astral^™^ mass spectrometer, depending on the batch, both equipped with an Easy Spray Source^™^ and a FAIMS Pro^™^ unit (following the configuration described in the Thermo Fisher Scientific technical note 002879).

Samples from all batches were processed on a Vanquish Neo equipped with an IonOpticks Aurora Ultimate XT analytical column (IonOpticks – AUR3-25075C18-XT) using the 50 Samples Per Day (50SPD) method from Thermo Fisher Scientific (technical note 002879).

Peptides are separated on a C18 reverse phase column warmed at 50^°^C using a total gradient of 25 min. The mobile phases A and B had the following composition: 0.1% Formic Acid for phase A and 80% Acetonitrile mixed with 0.1% Formic Acid for phase B. Both phases have been mixed during the gradient in the following steps:1-4%B for 0.1min, 4-12% B for 1.8min, 12%-22.5% B for 10.1min, 22.5%-40%B for 7.5 min, and 99%B for 5.5min for the column wash. The flow rate of the gradient was 200 nL/min, except for the first 1.9min, when it was 450 nL/min.

Data from batch 2 were collected in DIA mode on an Orbitrap Exploris 480 equipped with an Easy Spray Ion Source and a FAIMS Pro Duo Unit. The following source parameters have been used: ITT temperature of 275 ^°^C, FAIMS compensation voltage of -45, and a total carrier gas flow of 4L/min. The Full scan parameters were 120000 resolution, 400-800m/z of scan range, 45% RF lens, 300% of normalized AGD target, and Auto for the maximum injection time mode. The DIA scan parameters were 400-800 m/z precursor mass range, 60000 resolution, 50 m/z isolation window, 10 number of scan events, 28% HCD normalized collision energy, 120m/z as first mass, 60% RF lens, 1000% Normalized AGC target, and 118ms of max injection time.

Data from batch 1 and 3 were instead collected in DIA mode on an Orbitrap Astral, also equipped with an Easy Spray Ion Source and a FAIMS Pro Duo Unit. The following source parameters have been used: ITT temperature of 275 ^°^C, FAIMS compensation voltage of -50, and a total carrier gas flow of 4 L/min. The Full scan parameters were 240000 resolution, 400-800m/z of scan range, 45% RF lens, 500% of normalized AGD target, and 100ms maximum injection time. The DIA scan parameters were 400-800 m/z precursor mass range, 80000 resolution, auto for the DIA window type, window placement optimization ON, DIA window mode m/z Range, 20 m/z isolation window, 25% HCD collision energy, 150-2000 m/z scan range, 45% RF lens, 800% Normalized AGC target and 40ms of max injection time.

Raw MS data were searched using Spectronaut v19.4^21^ against the reviewed human UniProt reference proteome (UP000005640), supplemented with common contaminants. Search parameters were set to directDIA default settings: mass tolerances for precursors and fragments were determined automatically based on dynamic calibration; carbamidomethylation of cysteine was set as a fixed modification, while oxidation of methionine and N-terminal acetylation were included as variable modifications. False discovery rate (FDR) thresholds were strictly controlled at 1% at both the precursor and protein levels.

### Transcriptomic library preparation and sequencing

Eluted mRNA was processed using the SMART-seq mRNA Single Cell LP kit with the Unique Dual Index kit (1-96) per the manufacturer’s instructions ^22^. Libraries were sequenced on an Illumina NovaSeq X Plus platform.

### Transcriptomic data analysis

Raw reads were aligned to the human reference genome GRCh38 using STAR. Gene-level counts were quantified using STAR. Downstream analysis was performed in R (v4.3) using Seurat (v5.3) ^23^. Transcriptomic data were normalized using SCTransform, and dimensionality reduction and unsupervised clustering were performed using PCA followed by UMAP on the first 30 principal components. Query cells were projected onto the Liu et al. ^17^ iBlastoid reference atlas using the Seurat MapQuery framework with FindTransferAnchors and SCTransform-normalized data for anchor identification. Predicted cell type labels and reference UMAP coordinates were extracted for each query cell. For supplementary pairwise single-cell comparisons, expression differences between individual cell pairs were quantified as the delta between normalized values, with thresholds of |Δ| > 1 for RNA (SCTransform-normalized) and |Δ| > 0.5 for protein (log-normalized). Cells were classified as higher in cell A, higher in cell B, or similar based on these thresholds. All visualizations, including lineage-specific density plots and multi-panel figures, were generated in R (v4.3+) using ggplot2 ^24, 25^ and patchwork ^26^.

### RNA–protein concordance analysis

Protein identifiers were mapped to gene symbols to harmonize the feature space between modalities. Protein groups annotated with multiple gene symbols were collapsed to the first listed symbol prior to feature matching across modalities. Correlation analysis was restricted to features detected in both modalities within each cell, and only cells with at least 150 paired features were included. Missing values were treated as zero to indicate non-detection, and features with zero variance within the analyzed subset were assigned a z-score of zero and did not contribute to downstream correlation estimates. Spearman rank correlation coefficients were calculated per cell between SCTransform-normalized RNA values and log-normalized protein values. To correct for technical depth, expression values for each feature were residualized against the number of detected genes per cell (n_Genes_RNA_) and the number of identified proteins per cell (n_Proteins_ID_) using a linear regression model:

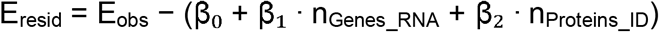

where E_obs_ is the observed expression level, and E_resid_ is the depth-independent residual. Partial Spearman ρ was subsequently calculated as the Pearson correlation of the ranks of these residuals.

### Differential expression and functional enrichment analysis

Differential expression analysis was performed using the MAST algorithm ^27^ on both SCTransform-normalized RNA and log-normalized protein data. Populations were defined by the top 10% of cells expressing SOX2 or GATA2. Significance thresholds were set at |log_2_FC| > 0.25 and unadjusted P < 0.05. For features detected in both populations, GO biological process enrichment was performed on the set of significantly differentially expressed features. For features detected exclusively in one population and absent in the other, GO enrichment was performed directly on the detected-only gene or protein set. Both analyses used clusterProfiler and org.Hs.eg.db (Bioconductor 2.23, 10.18129/B9.bioc.org.Hs.eg.db) with gene symbols mapped to Entrez IDs, and significant terms were defined by a Benjamini–Hochberg adjusted P-value < 0.05. Batch effects between the Astral and Exploris 480 datasets were corrected using the Harmony package (1.2.4) in R ^28^.

### Multi-omics factor analysis

Integrated analysis of transcriptomic and proteomic data was performed using the MOFA2 R package ^29^. A model was trained on 10 factors across the RNA and protein views. Factor scores (Z) and feature weights (W) were extracted for each factor. Variance explained (R^2^) was calculated for each factor and view. To assess cross-modal coupling, signed signature scores were computed per cell as the difference between the mean z-scored expression of the top 50 positive- and top 50 negative-weighted features in each view. Coupling was quantified as the Spearman correlation between factor scores and view-specific signature scores. Associations between factor scores and lineage identity were evaluated using a two-sided Wilcoxon rank-sum test with Benjamini–Hochberg FDR correction. Effect sizes were estimated using Cliff’s Delta. Over-representation analysis of top factor features was performed against the GO Biological Process database using ClusterProfiler ^30^, with significance defined at FDR < 0.05. All figures were generated in R (v4.3) using ggplot2 ^25^ and patchwork ^26^.

## Data availability

Mass spectrometry data are available via ProteomeXchange with identifier PXD076817. Raw RNA sequencing data have been deposited in the Gene Expression Omnibus (GEO) under accession number GSE327626. Analysis code and processed data tables used to generate all figures are available at https://github.com/DeMarcoLab/SC-multiomics. All other data supporting the findings of this study are available from the corresponding author upon reasonable request.

## Contribution

XX, MPC, SML, SAB, GS, NH, RV performed the experiments, XX, MPC, MLW, DHM, JP, RS, AdM analyzed the data, XX, AdM, and RS conceived the experiments. XX, MPC, MLW, and AdM wrote the article. All authors contributed to final editing and verified the information in the manuscript This work was supported by the NHMRC grant 2001066 and by the Simons Foundation SRAMM program.

