## Supplementary Figures and Tables for "Simultaneous single-cell profiling of the transcriptome and proteome"

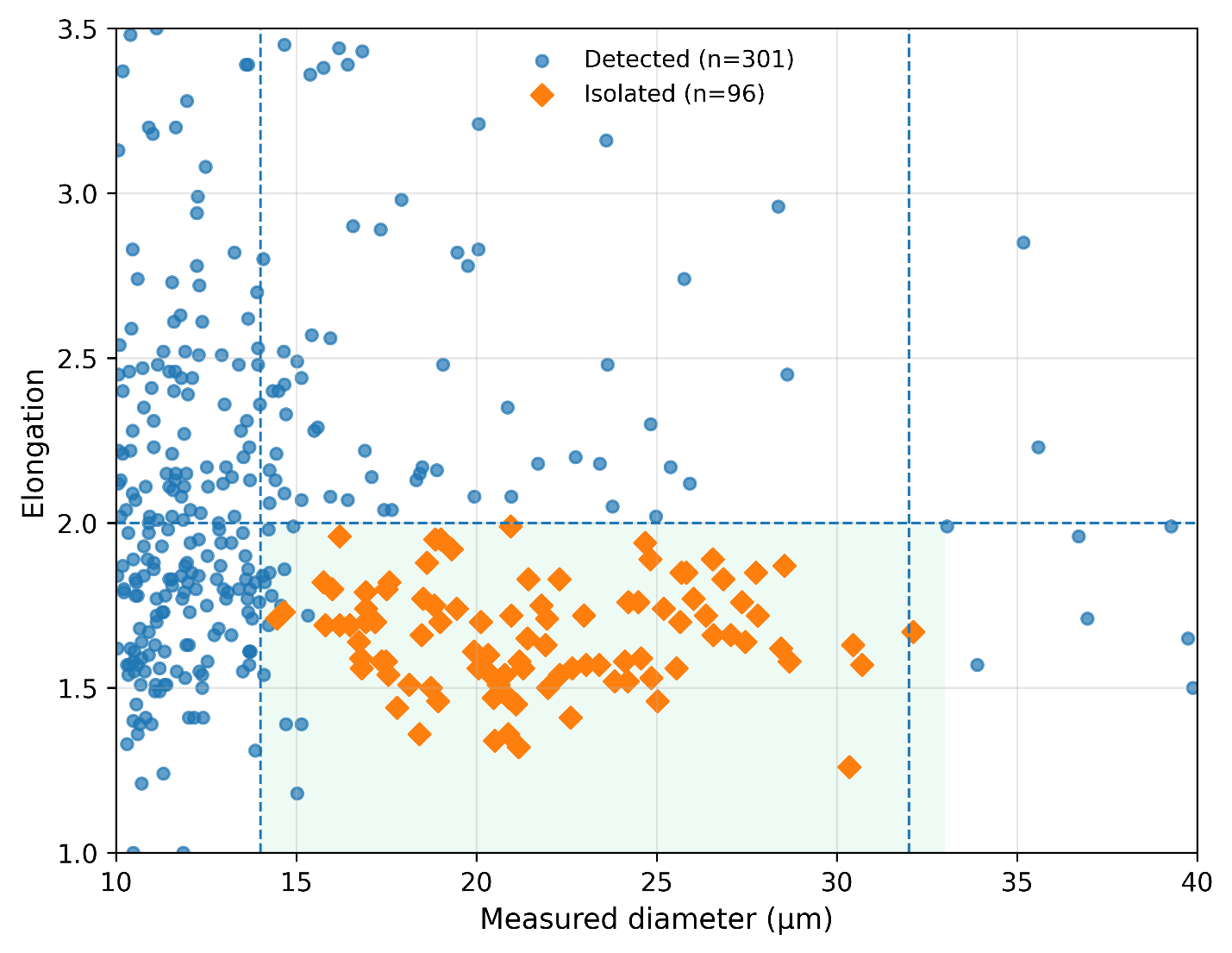

**Supplementary Figure 1 | Image-based selection of single cells using CellenOne.** Scatter plot showing measured cell diameter and elongation for all detected events and isolated single cells across three independent runs. Each point represents an individual particle identified by the cellenONE system. The shaded region indicates the pre-defined selection gate (diameter range of 14–32 µm and elongation < 2) used during dispensing. Dashed lines mark the corresponding threshold boundaries. In total 397 single particles were detected during sorting, of which 96 met the selection threshold and were isolated.

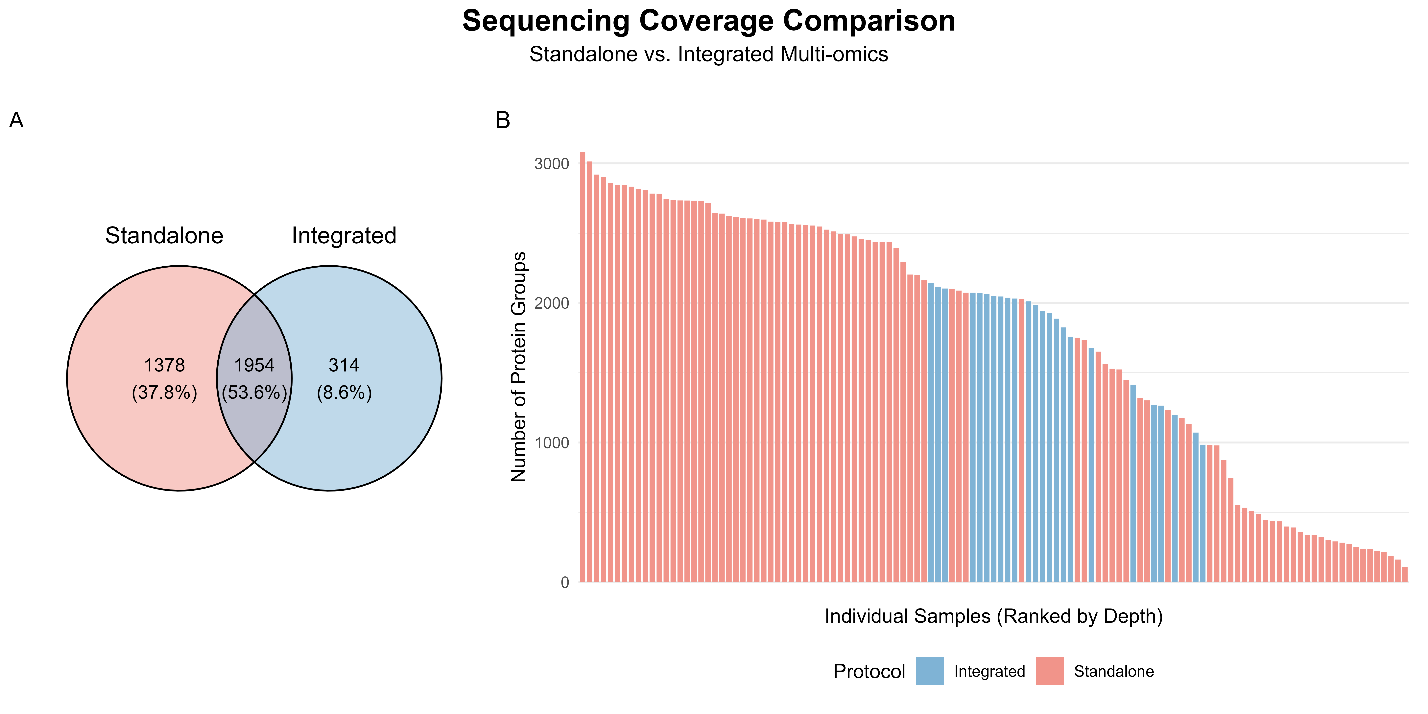

**Supplementary Figure 2 | Comparison of protein identification between standalone single-cell proteomics and integrated multi-omics workflows.** (A) Venn diagram showing the overlap of protein groups identified across all cells between the standalone and integrated workflows. A total of 1,954 protein groups (53.6%) were commonly detected by both approaches. In contrast, 1,378 protein groups (37.8%) were uniquely identified in the standalone workflow, while 314 protein groups (8.6%) were unique to the integrated workflow. These results indicate a substantial shared proteome coverage, with additional depth achieved in the standalone condition. (B) Bar plot showing the number of protein groups identified per individual cell, ranked by proteome depth. Each bar represents a single cell, with colors indicating workflow (standalone: red; integrated: blue). The integrated workflow yielded a mean of approximately 1790 protein groups per cell compared with approximately 1775.3 for the standalone protocol. Median was 1964.5 for the integrated workflow, compared with 2204 for standalone protocol. Despite this reduction, the overall distribution remains comparable between workflows, indicating that incorporation of the mRNA extraction step preserves a substantial proportion of proteome coverage at the single-cell level.

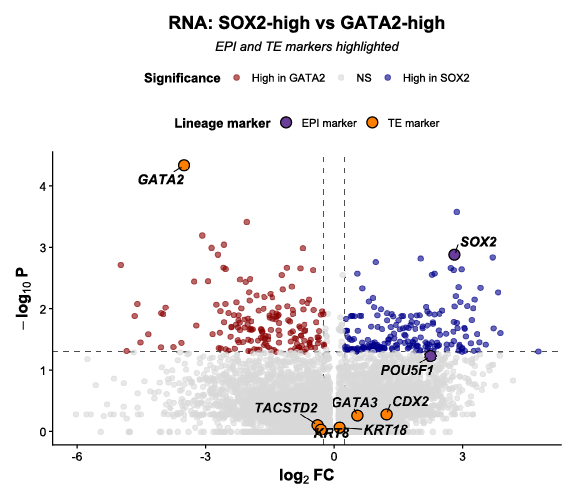

**Supplementary Figure 3 | Volcano plots RNA expression in cells displaying high GATA2 and high SOX2 markers.** This is the same plot as in Figure 2B, with the gene markers for the EPI and TE cell lineages highlighted.

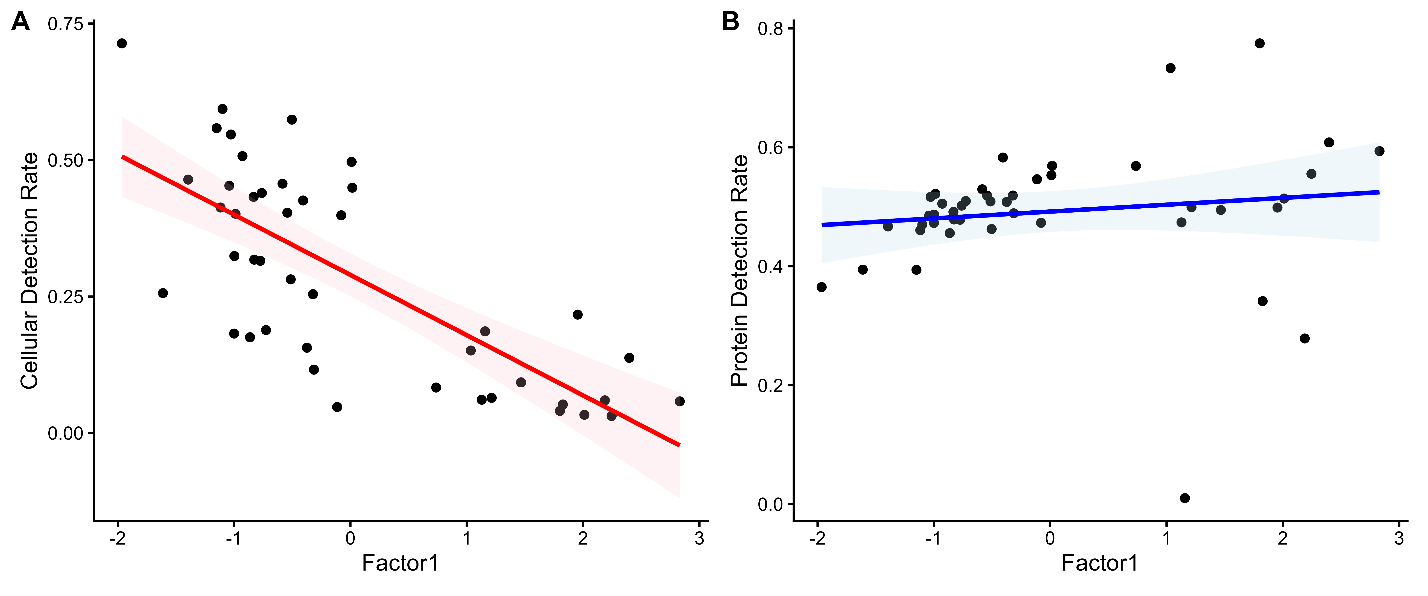

**Supplementary Figure 4 | Association of Factor 1 with cellular and protein detection rates.** (A) Scatter plot showing the relationship between Factor 1 values and cellular detection rate (CDR). Each point represents a single cell. A strong negative correlation is observed (r = −0.74), indicating that higher Factor 1 values are associated with reduced RNA detection per cell. (B) Scatter plot showing the relationship between Factor 1 values and protein detection rate (PDR). In contrast to CDR, only a weak positive association is observed (r = 0.13), suggesting that Factor 1 has some, but limited influence on protein detection depth.

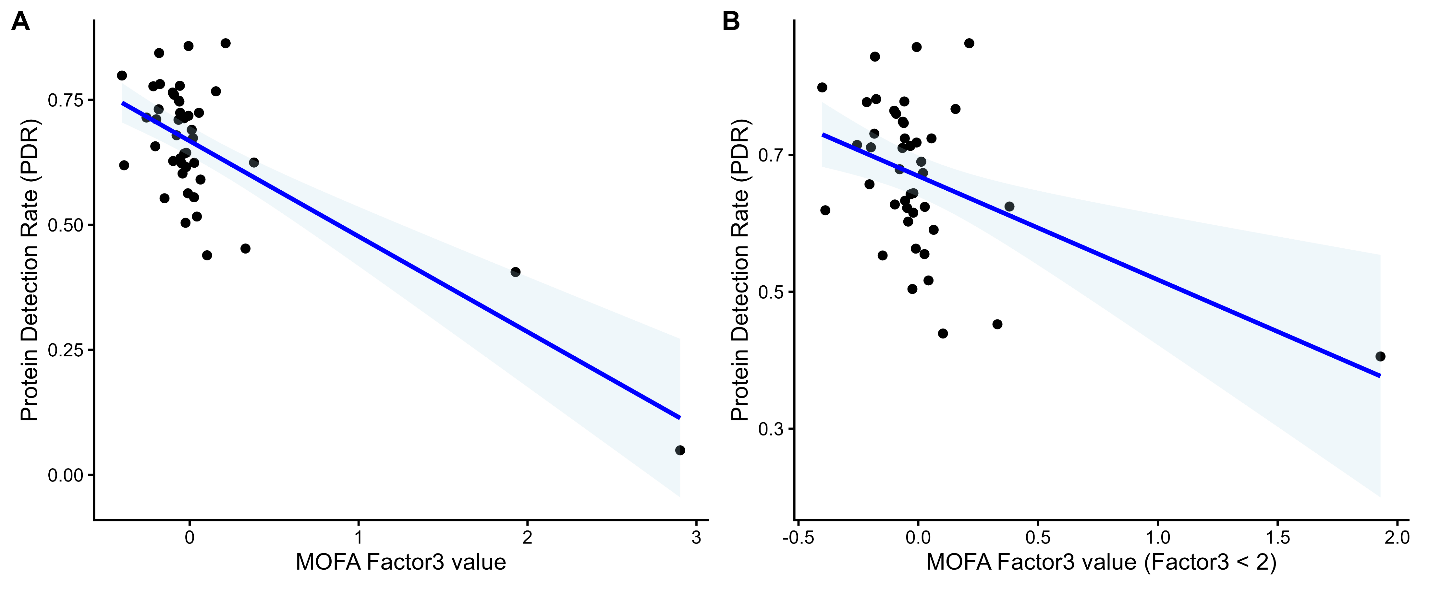

**Supplementary Figure 5 | Association of Factor 3 with protein detection rate.** (A) Scatter plot showing the relationship between Factor 3 values and protein detection rate (PDR) across all cells. Each point represents a single cell. A strong negative correlation is observed (r = −0.78), indicating that higher Factor 3 values are associated with reduced protein detection depth. (B) Scatter plot showing the same relationship after excluding extreme values (Factor 3 < 2). The negative association remains strong but is attenuated (r = −0.56), suggesting that factor 3 strongly correlates to PDR.

**Supplementary Figure 6 | Gene ontology enrichment of top loading features for Factor 2.** Dot plot showing Gene Ontology (GO) enrichment analysis of the top loading features associated with Factor 2 in the MOFA2 model. Each dot represents a significantly enriched GO term, with size indicating the number of contributing features and color representing the adjusted p-value. The x-axis denotes the gene ratio for each term. Enriched terms are predominantly related to ribonucleoprotein complex biogenesis, membrane-less organelle assembly, and stress granule formation. Additional terms include regulation of cell fate specification and protein export from the nucleus. These results indicate that Factor 2 is associated with variation in ribosome-related processes and post-transcriptional regulation, consistent with its strong contribution from the proteomic modality.

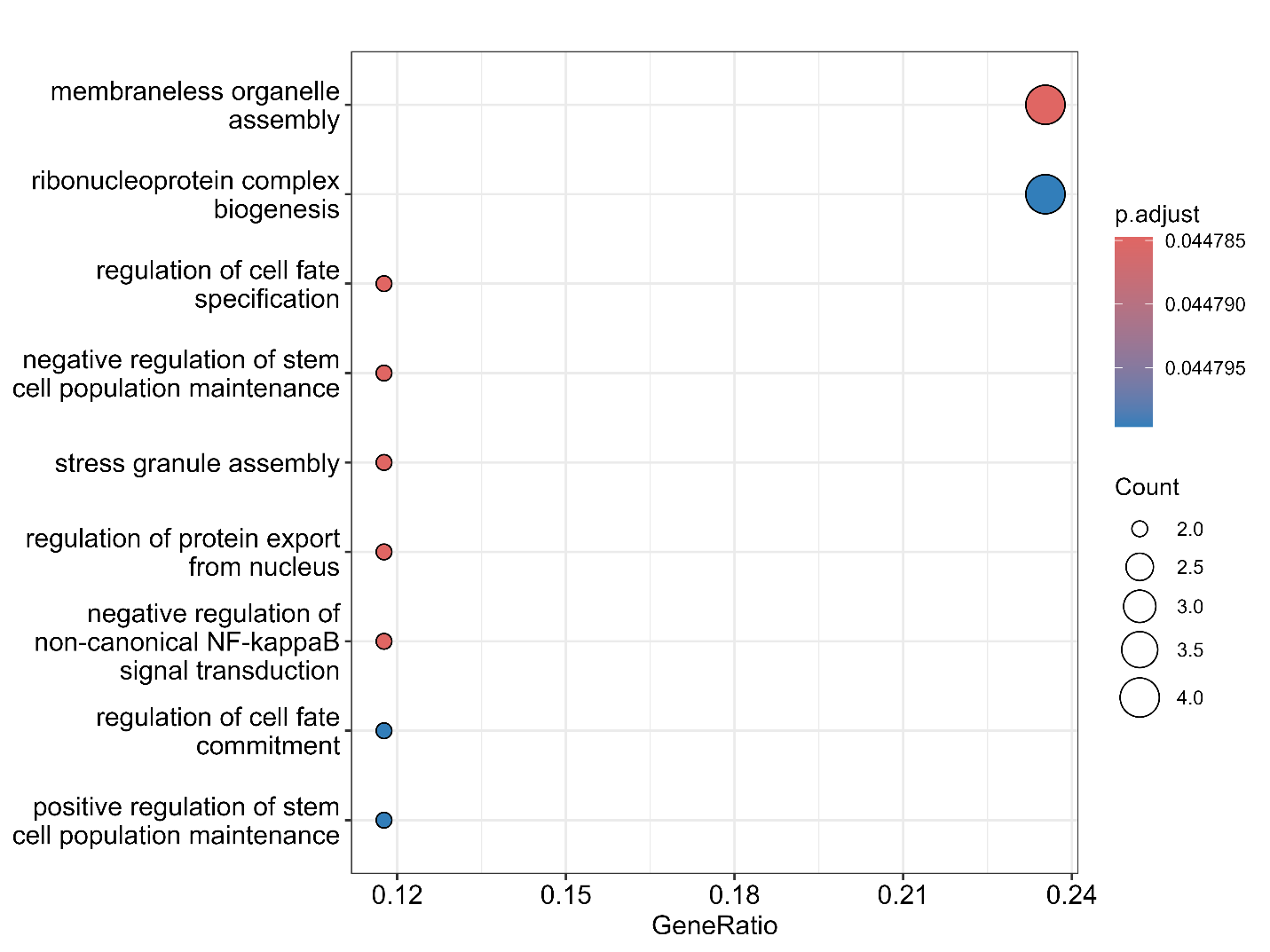

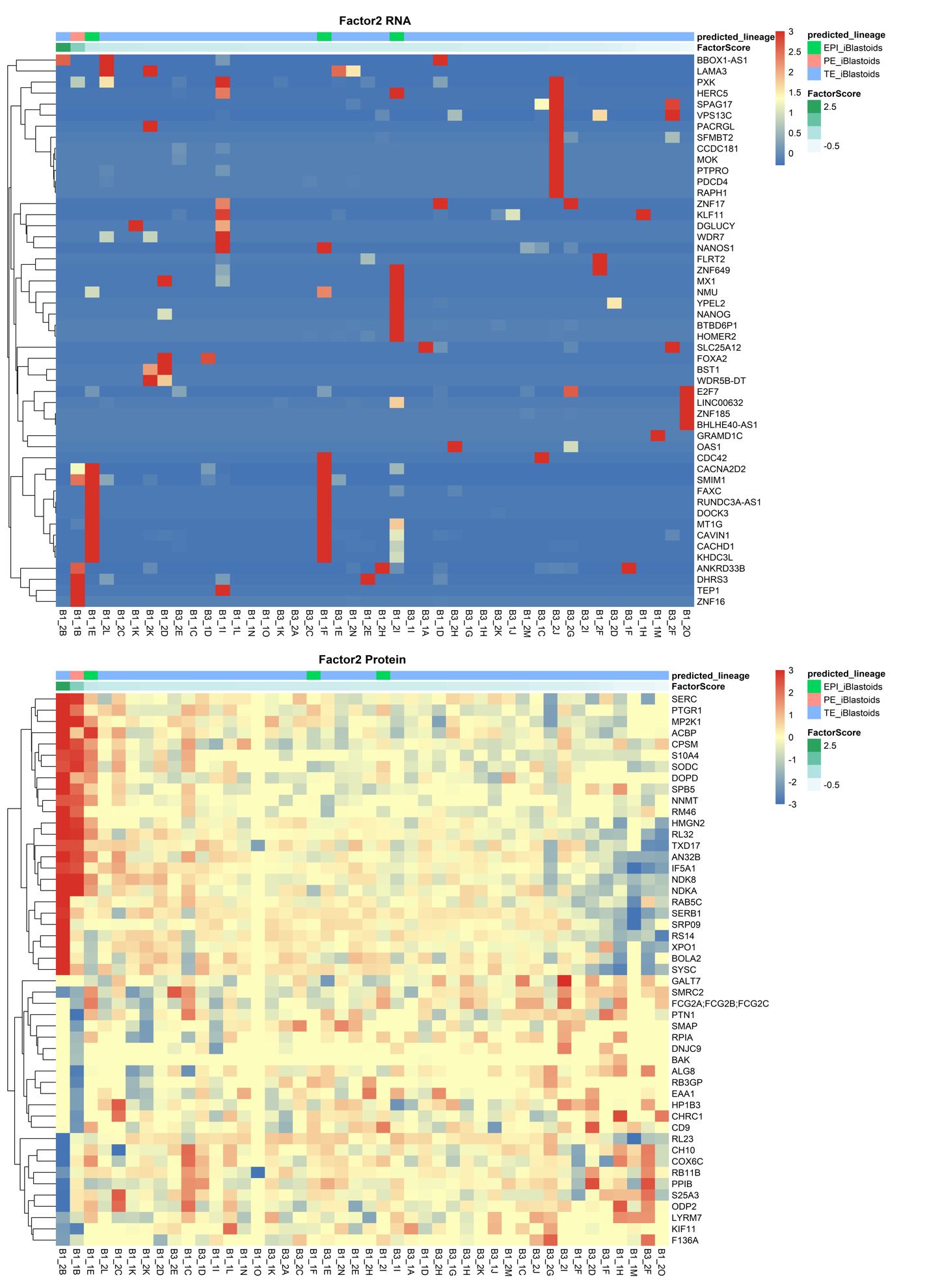

**Supplementary Figure 7 | RNA and protein feature loadings associated with Factor 2.** Heatmaps showing the top loading features for Factor 2 across RNA (top) and protein (bottom) modalities. Cells are ordered by Factor 2 values, and features are selected based on the highest absolute loadings in each modality. Expression values are row-scaled (z-score) and capped to improve visual contrast, as implemented in the analysis pipeline. For each modality, the heatmaps include both positive and negative loading features. The top 25 rows correspond to features with the highest positive loadings, while the bottom 25 rows correspond to features with the most negative loadings. This layout highlights opposing patterns of variation associated with Factor 2 across cells.

The RNA heatmap shows sparse and heterogeneous patterns with limited structure across the Factor 2 axis. In contrast, the protein heatmap displays more coherent and coordinated variation, with distinct gradients corresponding to Factor 2 values. Annotation bars indicate predicted lineage and Factor 2 scores for each cell.

**Supplementary Figure 8 | Association of Factor 4 with lineage identity and technical covariates.** (A) Distribution of Factor 4 values across predicted lineage groups. Factor 4 shows a significant association with lineage identity (Kruskal–Wallis p = 0.0147). Pairwise comparison confirms a significant separation between TE-like and EPI-like populations (BH-adjusted p = 9.7 × 10⁻⁴). Lineage identity explains a substantial proportion of variance in Factor 4 (R² = 0.63), indicating strong biological relevance. (B) Scatter plot showing the relationship between Factor 4 values and cellular detection rate (CDR). Only a weak positive correlation is observed (r ≈ 0.11), suggesting minimal influence of RNA detection efficiency on Factor 4. (C) Scatter plot showing the relationship between Factor 4 values and protein detection rate (PDR). The association remains weak (r ≈ 0.07), indicating limited contribution from protein detection depth.
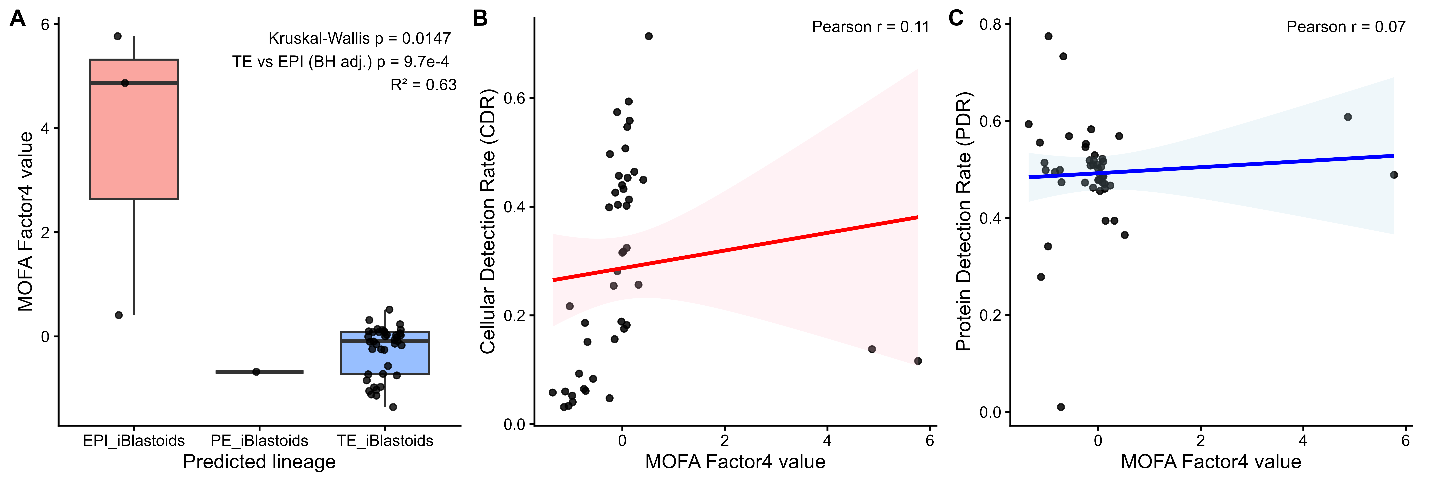

**Supplementary Figure 9 | Relationship between Factor 2 and Factor 4 colored by predicted lineage.** Scatter plot showing single cells projected by MOFA Factor 2 and MOFA Factor 4, with points coloured by predicted lineage. Factor 2, interpreted as a biologically informative factor, is plotted against Factor 4 to assess whether Factor 4 aligns with lineage structure in the multi-omic dataset. The plot was generated directly from MOFA factor values and lineage metadata, with Factor 2 and Factor 4 extracted per cell and colored by the predicted_lineage annotation. Cells assigned to the EPI-like lineage occupy a distinct region with high Factor 2 values, whereas TE-like cells cluster near lower Factor 2 values and show broader spread along the Factor 4 axis. This separation supports the interpretation that Factor 4 is associated with lineage identity rather than random technical variation. In this context, plotting Factor 4 against another biologically relevant factor provides an orthogonal view of lineage structure and highlights the clear relationship between Factor 4 and predicted cell state.
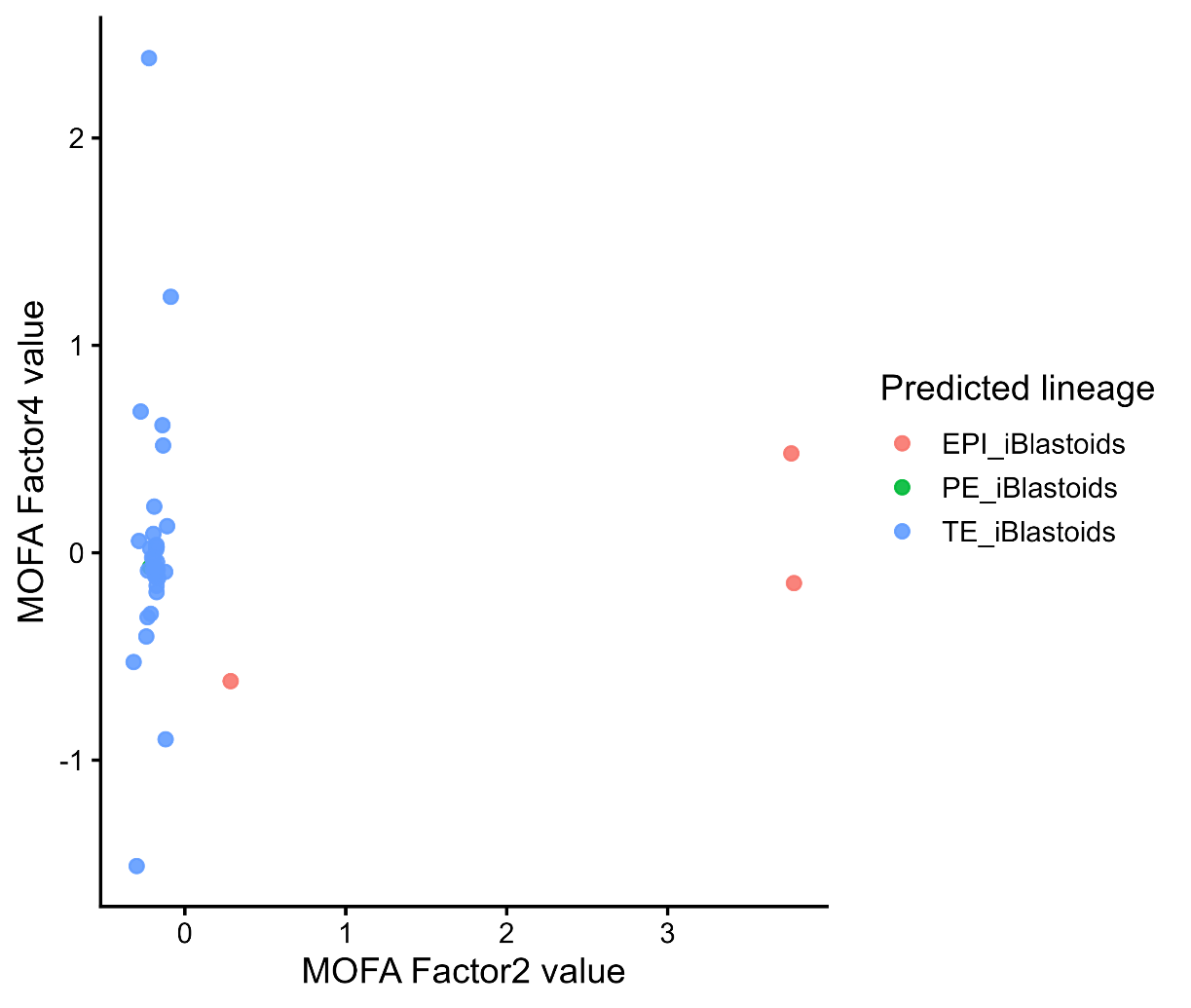

**Supplementary Figure 10 | Top RNA features associated with Factor 4 in the MOFA2 model.** Scatter plots showing the relationship between MOFA Factor 4 values and the expression of the top 10 RNA features selected by weight in the RNA view. Each panel corresponds to one feature identified by the MOFA2 plot_data_scatter function, and points are colored by predicted lineage. A fitted linear regression line is shown in each panel together with the corresponding correlation coefficient and p-value. Most top-weighted RNA features show strong positive associations with Factor 4, with correlation coefficients ranging from 0.81 to 0.93. Cells with higher Factor 4 values are predominantly assigned to the EPI-like lineage, whereas TE-like cells cluster at lower Factor 4 values. This pattern is consistent across multiple genes, indicating that Factor 4 is linked to a coherent lineage-associated transcriptional program rather than isolated feature-specific effects.

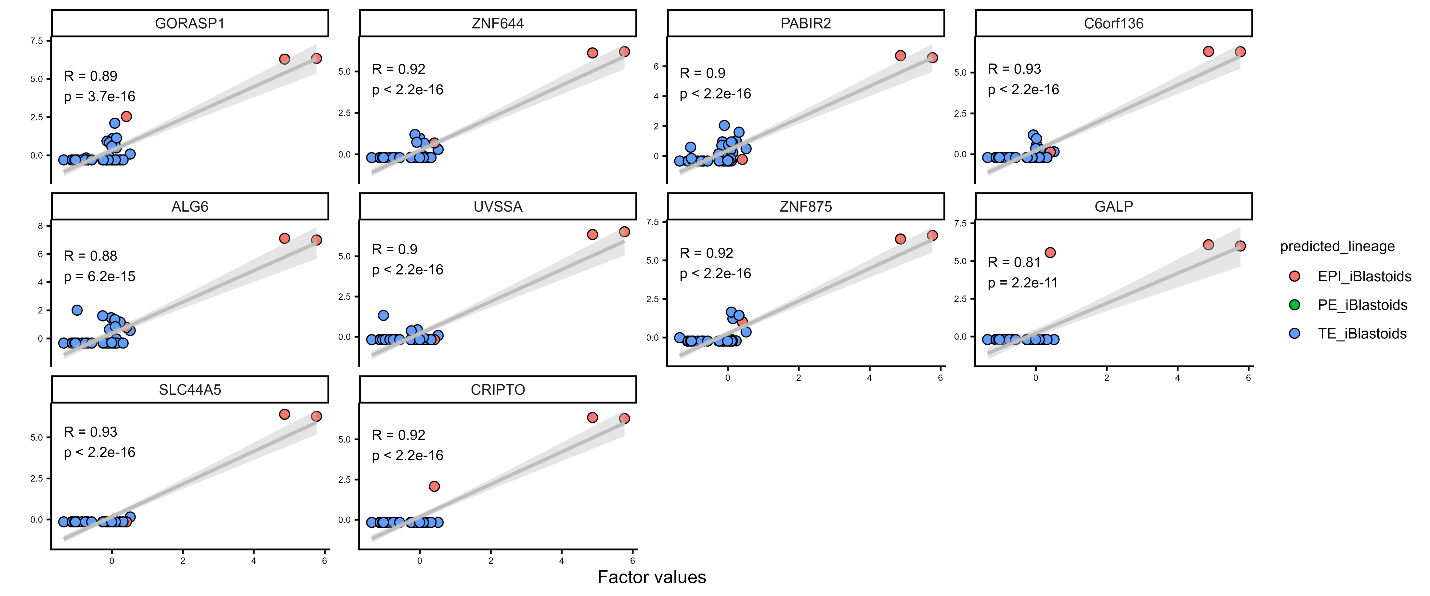

| **Factor** | **RNA**  **Variance Explained** | **Protein**  **Variance Explained** | **Dominant Modality** | **Biology** | **Factor Type** | **Lineage Association** | **QC Association** |
| --- | --- | --- | --- | --- | --- | --- | --- |
| F1 | High | Low | RNA | Global transcript detection | Global technical | No | Strong (CDR) |
| F2 | Low | ~14% | Protein | RNP biogenesis / translation activity | Biological state | No | Minimal |
| F3 | ~0 | Moderate | Protein | Proteome detection rate | Technical (protein-specific) | No | Strong (PDR) |
| F4 | Moderate | Moderate | Shared | Lineage axis (TE vs EPI) | Biological lineage | Yes (R²=0.63) | Minimal |
| F5 | ~0 | ~6.5% | Protein | Mitochondrial / OXPHOS metabolic program | Biological metabolic state | Weak | Minimal |
| F6 | ~0 | ~4% | Protein | Keratin / desmosome structural module | Protein structural module | No | Minimal |
| F7 | ~0 | ~3% | Protein | Metabolism / redox | State / partial QC mix | No | Partial (PDR) |
| F8 | ~0 | ~2.7% | Protein | Mitochondrial vs nuclear growth allocation | Biological state | No | Low |
| F9 | ~0 | ~2% | Protein | Mitochondrial /proteostasis–RNP gradient | QC-state axis | No | Minimal |
| F10 | ~0 | ~1.8% | Protein | Nuclear RNP/DNA repair vs chaperonin axis | Protein state | No | Minimal |

**Supplementary Table 1 | Overview of factors identified in Multi-Omics Factor Analysis**. Multi-Omics Factor Analysis revealed 14 factors in the multi-omic dataset. While the initial model was parameterized for 10 factors to capture the primary axes of variation, the final trained model converged on 14 factors, indicating additional subtle but consistent sources of variation. Factor 1 to 10 had at least one view with variance explained (R^2^)> 1% and were considered for downstream analysis.
